# A3D 2.0 update for the prediction and optimization of protein solubility

**DOI:** 10.1101/2021.04.13.439600

**Authors:** Jordi Pujols, Valentín Iglesias, Jaime Santos, Aleksander Kuriata, Sebastian Kmiecik, Salvador Ventura

**Author notes:** To whom correspondence should be addressed: Salvador Ventura, Institut de Biotecnologia i Biomedicina, Parc de Recerca UAB, Mòdul B, Universitat Autònoma de Barcelona, E-08193 Bellaterra (Barcelona).

## Abstract

Protein aggregation propensity is a property imprinted in protein sequences and structures, being associated with the onset of human diseases and limiting the implementation of protein-based biotherapies. Computational approaches stand as cost-effective alternatives for reducing protein aggregation and increasing protein solubility. AGGRESCAN 3D (A3D) is a structure-based predictor of aggregation that takes into account the conformational context of a protein, aiming to identify aggregation-prone regions exposed in protein surfaces. Here we inspect the updated 2.0 version of the algorithm, which extends the application of A3D to previously inaccessible proteins and incorporates new modules to assist protein redesign. Among these features, the new server includes stability calculations and the possibility to optimize protein solubility using an experimentally validated computational pipeline. Finally, we employ defined examples to navigate the A3D RESTful service, a routine to handle extensive protein collections. Altogether, this work is conceived to train and assist A3D non-experts in the study of aggregation-prone regions and protein solubility redesign.

## 1. Introduction

Protein aggregation is a deleterious process that leads to the formation of protein deposits. Despite being classically associated with disease-related proteins, aggregation is a generic trait of polypeptide chains, being conserved across proteomes despite millions of years of evolution (1). The ubiquitous presence of aggregation determinants in proteins is the consequence of the molecular determinants governing aggregation overlapping with the physicochemical properties required to establish native contacts and perform essential functions (2–4). Accordingly, proteins have evolved under the permanent und unavoidable pressure of aggregation, adjusting aggregation loads to the necessary minimum required for function and solubility (5).

In vivo, protein solubility and aggregation are in a constant equilibrium that is surveilled by the protein quality control machinery (6). However, specific scenarios such as genetic mutations, abnormal post-translational modifications, or the deterioration of proteostasis networks may push proteins out of their natural context and favor nucleation reactions and, ultimately, aggregation (7). Similarly, along the pipelines of biotechnological processes, proteins are exposed to non-native buffering conditions and concentrations that exceed the cellular levels by several orders of magnitude (8). Proteins have not evolved to remain soluble in these scenarios, and accordingly, they precipitate in cells in human diseases or during the production processes of therapeutic proteins. In this context, there is an urgent need to develop straightforward approaches to predict protein solubility and identify optimized variants. Protein aggregation is usually triggered by the transient or permanent exposure of short polypeptide stretches with significant aggregation propensities. These aggregation-prone regions (APRs) are usually enriched in hydrophobic residues and are both sufficient and necessary to establish the non-native intermolecular contacts that nucleate aberrant self-assembly (9, 10). Accordingly, the amino acids’ physicochemical properties, together with their specific position in the polypeptide chain, determine the aggregation propensity of proteins. The structural context of folded proteins, including their tertiary and quaternary structures, modulates this aggregation load. Spatial clustering of sequentially nonconsecutive hydrophobic residues can generate Structural aggregation-prone regions (STAP), which recapitulate the properties of sequential APRs in protein surfaces.

The implementation of computation algorithms that exploit our theoretical and experimental knowledge about protein sequences and structures has become a cost-effective approach to understand and manipulate aggregation (11). More than 30 predictive algorithms have been designed to identify both APRs and STAPs in protein sequences and structures (12), offering a versatile toolbox to assess protein aggregation from different perspectives and serve the scientific community with an appropriate pipeline for each specific study case. These computational tools have demonstrated their efficacy not only in predicting individual protein aggregation but also in anticipating the impact of mutations, performing proteome-wide analysis, and assisting in the redesign of solubility of proteins with biotechnological interest (13). AGGRESCAN 3D (A3D) is our in-house structure-based algorithm to predict protein aggregation upon protein structures (14). It employs an experimentally derived scale of aggregation propensity for each natural amino acid (15). These intrinsic aggregation values were first implemented in the linear aggregation predictor AGGRESCAN to analyze protein sequences (16). In A3D, amino acid aggregation propensities are projected into the 3-dimensional structure of the protein and adjusted as a function of their relative exposure to solvent. Then, the algorithm considers a spherical area of 10 Å, centered in the Cα carbon of the considered amino acid, and computes the aggregational influence of other residues included in this sphere. The spatial proximity, intrinsic aggregation propensity, and relative exposure of these amino acids are integrated to provide a defined aggregation score. Accordingly, A3D identifies STAPs in protein surfaces, while discarding the contribution of hydrophobic residues that are buried in protein cores. A3D also incorporates dynamic simulations of protein structures in its pipeline, which allows us to approximate the structural fluctuations of native conformations and uncover transiently exposed STAPs. Finally, it offers the possibility to mutate specific residues virtually and, therefore, predict the impact of solubilizing or pro-aggregation substitutions.

Since A3D’s launching in 2015, a significant number of studies have exploited this approach to enhance protein solubility (17–19). However, the original A3D version (14) had several limitations, and some helpful features were missing: i) dynamic runs were not allowed for proteins larger than 400 amino acids, having more than one polypeptide chain, or missing residues; ii) stability analysis upon mutation was not possible (overseeing thermodynamically unfavorable mutations leading to overall destabilization of the protein structure); iii) a significant knowledge on protein folding and protein aggregation principles was required to optimize protein solubility successfully; iv) large-scale predictions were restricted by the manual introduction and retrieving of protein structures, a time-consuming task that ultimately prevented including A3D in automated pipelines. We have recently updated A3D to a 2.0 version that overcomes these limitations and facilitates the A3D use for new users (20). In this work, we will illustrate these implementations and how to use them correctly.

## 2. Aggrescan 3D 2.0

The 2.0 version of A3D (20) is accessible online at http://biocomp.chem.uw.edu.pl/A3D2/. Server upgrade incorporates features addressing the limitations as mentioned earlier:

- The new stand-alone version of CABS-flex allows us to extend input entries for dynamic analysis to multi-chain structures, proteins up to 4000 residues, and incomplete PDBs (21).
- Protein stability, calculated using the FoldX force field, is now taken into account when analyzing the impact of protein mutations.
- The incorporation of an Automated Mutations widget generates structural variants with enhanced solubility and complements the manual mutation tool in the redesign of protein solubility.
- The visualization and managing of protein structures have been improved with a significantly faster display that includes different tools to inspect A3D 2.0 output structures.
- Implementation of a new RESTful service allows the use of A3D in high-throughput computational screenings or in-depth mutational analysis.

The implementation of these new characteristics expands the predictive potential of A3D to previously unsupported protein structures and integrates stability outputs into a more comprehensive and user-friendly interface for the redesign of protein solubility.

### 2.1. Front page

A3D 2.0 front page functions as the input interface of the server (Fig. 1). In the “*Input structure*” box, users must provide a valid PDB identifier or upload a PDB-formatted file from a local directory and indicate, if needed, the specific chain(s) that are expected to be analyzed; while no indication results in the analysis of all polypeptide chains. In the “*options*” box, users can set the parameters of the following run. The updated A3D 2.0 server keeps most of the utilities that were already available in the previous version of A3D (22). A3D 2.0 predictions can be run in static and dynamic modes. By default, A3D runs are executed in static mode, where no contributions of chain flexibility are considered. If needed, users can activate CABS-flex simulations on the prediction pipeline and run a dynamic analysis. Aggregation analysis can be performed with a 10 Å or 5 Å “*distance of aggregation analysis*, ” which directly impacts A3D’s calculations since it restricts the contribution of neighboring residues to the amino acid under evaluation. Accordingly, the aggregation scores of analyses performed with a 10 Å radius sphere will be influenced by more distant residues, and vice versa. By selecting the “*Mutate residues*” tool option, selected residues of input proteins can be manually mutated to one of the remaining nineteen amino acids. Finally, the “*options*” box also allows us to activate the new features “*enhance protein solubility*” and “*stability calculations*”.

**Figure 1.**
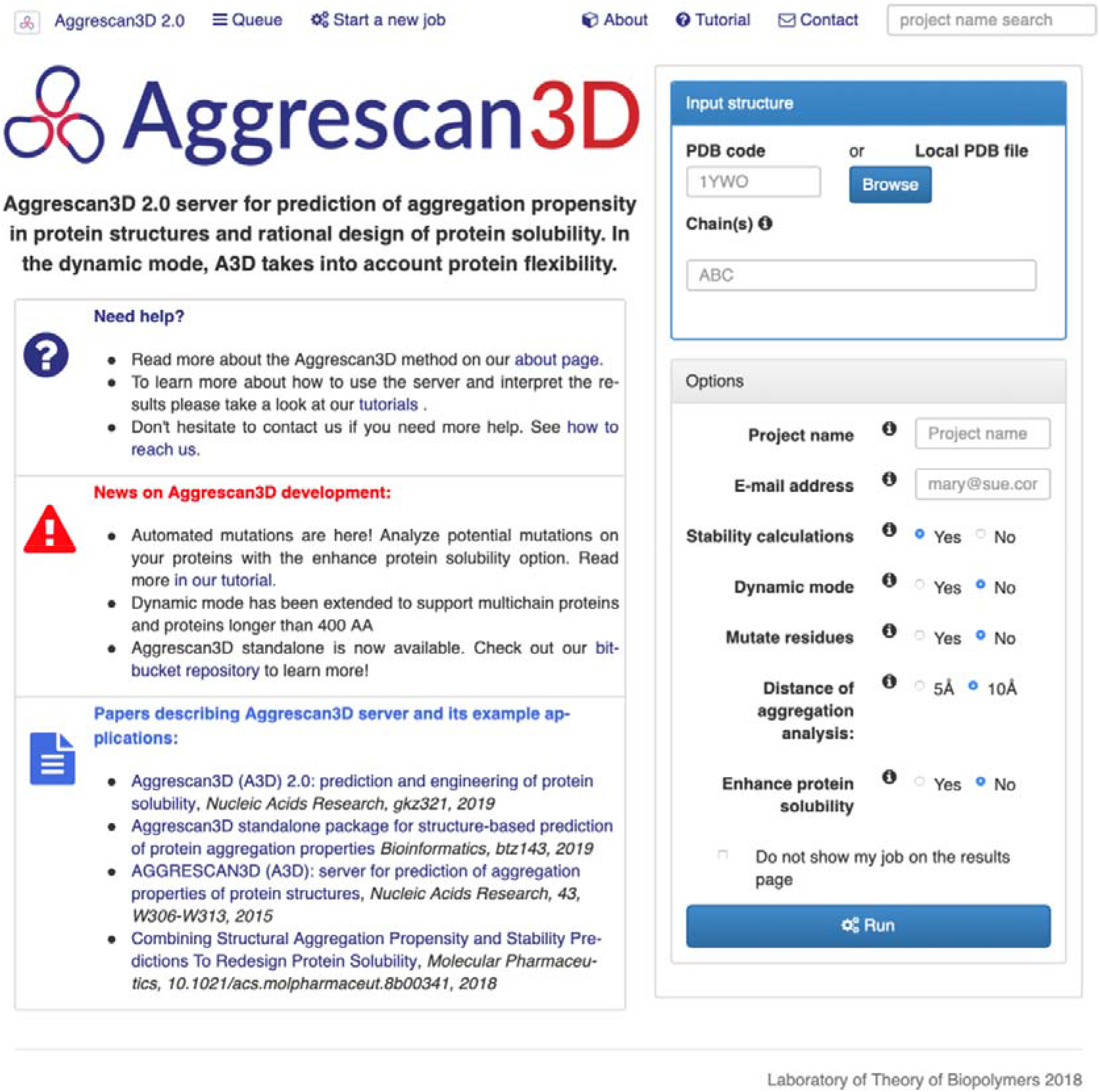
Frontpage of the updated A3D 2.0 server.

### 2.2. Stability calculations

In order to perform their biological function, globular proteins fold, and adopt well-defined three-dimensional structures. Functional conformations are thermodynamically stable and correspond to energetically favored states (23). However, sequence alterations, such as simple point mutations, insertions, or deletions, can destabilize protein structures and their complexes, associated with loss of function and aggregation events. In this context, one of the major concerns when designing protein variants with enhanced solubility by introducing mutations is their repercussion in the structural stability and, thus, in protein function (24–26). The updated A3D 2.0 analysis implements stability calculations to assist users’ endeavors in protein engineering and report on the thermodynamic contribution of the introduced mutations.

FoldX is an empirical force field that has been extensively used to model the stability of protein structures (27). It represents a valuable tool when assessing the effect of protein mutations because it computes the energetic difference between the *wild type* protein and its redesigned variant (*see* **Note** Error! Reference source not found.). A negative difference of free energy (ΔΔG<0) accounts for energetically favorable modifications, and a positive difference of free energy (ΔΔG>0) accounts for an energetically unfavorable mutation (*see* **Note 2**). Mutations with ΔΔG>0.5 kcal/mol are considered as significantly destabilizing (28).

In A3D 2.0, FoldX’s stability calculations are only operative if a mutation tool is activated. In the case of the classic manual mutation tool, free energy calculations are displayed in the “*project details*” tab of completed jobs. For the new automatic mutations tool “enhanced solubility”, the information referring to stability is displayed in the “*automated mutations”* output tab (see enhanced solubility section for more information).

### 2.3. Dynamic calculations

A distinctive feature of A3D is the possibility of using dynamic simulations of protein structures to predict aggregation. Despite being compact folded structures, most of the proteins usually contain regions with certain flexibility (29). This structural dynamism is associated with the attainment of protein interactions, which are considered a driving force for protein function (30). As a consequence of structural rearrangements, proteins might expose sheltered regions with significant hydrophobicity (31). These APRs would be classified as non-exposed residues and therefore considered as non-contributing regions in static runs, although their temporary exposition may be enough to initiate the aggregation cascade (32). The A3D 2.0 dynamic approach simulates fluctuations of protein structures in solution and approximates the aggregation behavior of proteins in their natural environments.

The A3D 2.0 pipeline integrates the new stand-alone application version of CABS-flex (21) for fast structure flexibility simulations. CABS-flex is based on coarse-grained modeling of polypeptide chains, which significantly shortens simulation times compared to all-atom molecular dynamics (21). In the stand-alone application version, CABS-flex allows the simulation of multi-chain structures up to thousands of amino acids. Therefore, now, the applicability of the A3D 2.0 dynamic mode extends to large biologically relevant proteins such as antibodies, protein oligomers, multimeric enzymes, and heteromeric protein complexes (*see* **Note 3**). When activated, the dynamic mode pipeline uses the CABS-flex method to generate 12 structural models (representative of the conformational states generated in the CABS-flex simulations) and compute aggregation propensities for all of them. Accordingly, users get a spectrum of aggregation tendencies according to different conformational states around the initial structure. Run results are shown in the *“dynamic mode details”* tab of completed jobs (Fig 2). As an example, we analyze the amyloid core of the cytoplasmic polyadenylation element-binding of *Drosophila melanogaster,* Orb2, recently solved by Cryo-EM. Of note, Orb2 PDB comprises nine different chains, being inaccessible to A3D’s previous configuration. The average scores for all generated models (numbered from 0 to 11) and the input structure are displayed in the left table and ranked according to their aggregation propensity. By clicking the “*view”* button, users can visualize each model, colored with the classic A3D aggregation scale (see structure visualization and gallery section for more information). The three-dimensional coordinates for all protein models can be downloaded by clicking the “*pdb”* button. Likewise, “*CSV”* prompts the download of a comma-separated values (.csv) file containing the aggregation scores computed for individual residues. These aggregation scores are plotted as a function of the residue number in an interactive graph in the lower section of the results’ page (*see* **Note 4**). Users can select one or more models to be displayed just by clicking the specific model’s name on the graph’s legend. Moving the cursor through the plotline pops residue information up (chain, residue name, residue number, solubility prediction, and model). Finally, in the bottom section, the input structure (red) and the most aggregation-prone model (blue) are superimposed, and the root-mean-square fluctuation (RMSD) plotted as a proxy for local protein flexibility (*see* **Note 5**). All in all, these data grant the opportunity to compare in a fast and comprehensive interface the input structure with CABS-flex models and to identify specific regions that might link aggregation properties and structure flexibility (*see* **Note 6**).

**Figure 2.**
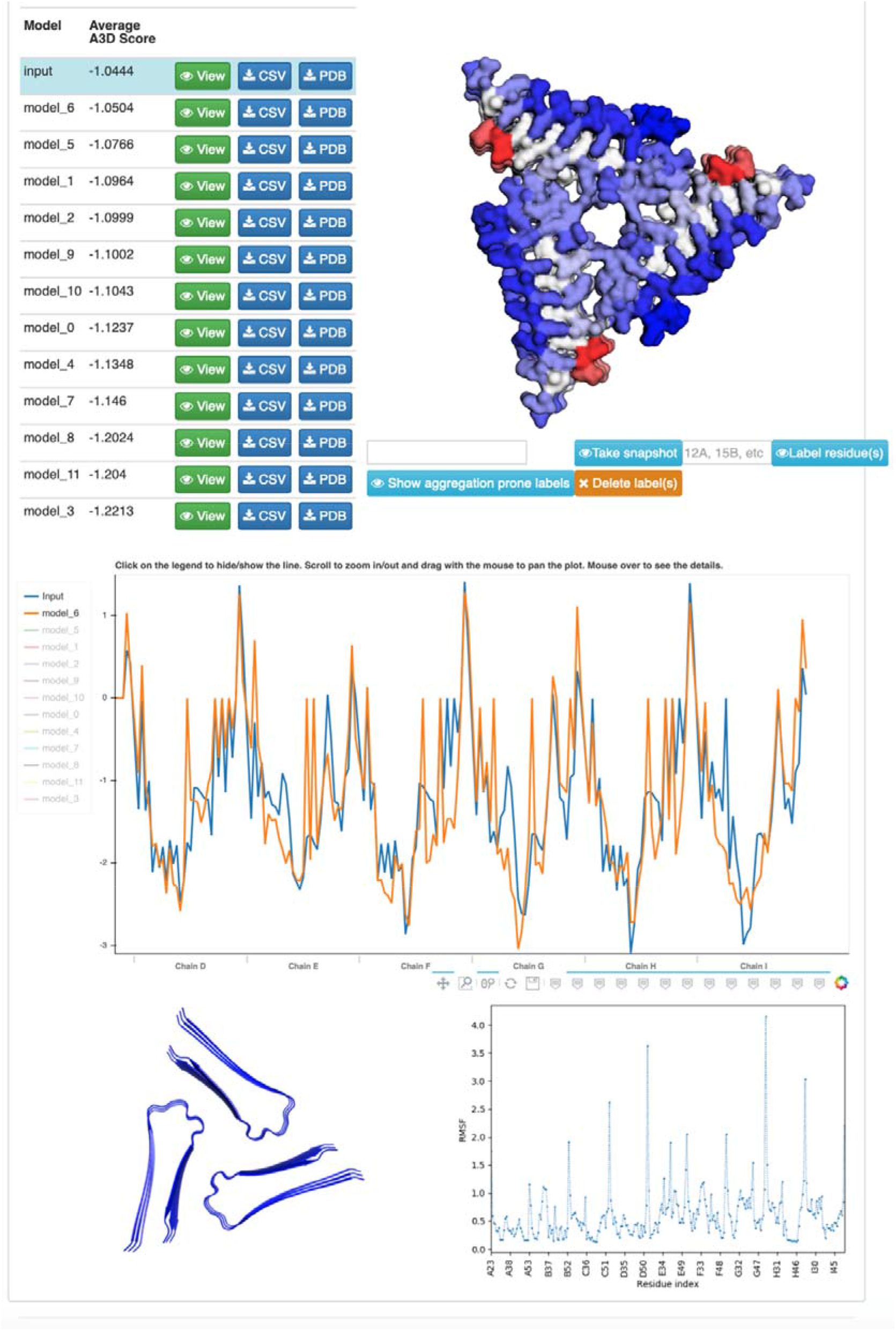
A3D 2.0 dynamic mode output for Orb2 protein (PDB: 6VPS). Predictions for the input structure and the 12 CABS-flex simulations are displayed in the table. A3D 2.0 incorporates an interactive interface to analyze each conformation with the structure visualization tool (top right) and the aggregation propensities plot (middle). In the bottom section, users can find information about the RMSD of the top-scoring CABS-flex model and its structural overlapping with the input structure.

### 2.4. Automated mutations tool

Protein aggregation is a significant drawback in the synthesis, storage, and administration of protein-based therapies (33, 34). Promising drugs such as antibodies and replacement enzymes can precipitate along the production pipeline, which dramatically decreases their purity and yield (35). Likewise, before reaching the eventual patient, they are usually conserved at highly concentrated stock solutions, where aggregation can occur (36). This eventual aggregation alters the activity of drug molecules and compromises patient’s health, triggering an immunogenic response against such aggregated species (37, 38). Overall, aggregation may prevent the commercialization of otherwise promising therapeutics.

To minimize the impact of aggregation, pharmaceutical companies dedicate considerable efforts to the redesign of protein variants with enhanced solubility, usually with the implementation of direct evolution methodologies, such as phage display (39). Computational predictors of protein aggregation stand as cost and time effective alternatives to these approaches, which, in the end, are based on blind combinatorial experiments (40). The novel automated mutations tool was implemented in A3D 2.0 to complement the already present manual mutation tool and facilitate protein engineering to non-expert users (*see* **Note 7**).

The automated mutations tool is accessible at the front page as the “*Enhance protein solubility*” option. The tool automatically identifies residues with the highest A3D score in an initial run in the static mode (*see* **Note 8**). Then, up to six most aggregation-prone residues (only those with positive A3D score) are individually mutated to positively (Lysine and Arginine) or negatively charged residues (Aspartic and Glutamic acids), thus computing four possible solubilizing variants for each single amino acid under consideration (i.e., 12 candidates in total in case of 3 residues) (*see* **Note 9**). Before starting the analysis, users are redirected to a new window, where the amino acids required for protein function can be excluded from the study (*see* **Note 10**). All the protein variants are re-analyzed in the static mode as independent structures and ranked as a function of A3D average aggregation score and stability increments, taking the input structure as a reference. Thermodynamically unfavorable (ΔΔG>0) and pro-aggregation (ΔA3D_score_>0) mutations are strongly penalized; thus, destabilizing mutations are computed as low-ranking candidates, even if they are also extremely solubilizing variants.

Accessible in the “*Automated mutations*” tab, the algorithm renders a limited list of top-ranked mutations (Fig 3). The displayed list includes two top-ranked mutants for each mutated residue, while the complete list containing all the mutants can be downloaded as a .csv file. Each displayed mutant structure can be visualized and explored in depth by clicking the following buttons: “*View*” (online display of a mutant structure that is colored according to A3D scale); “*CSV*” (download A3D scores for each residue of a mutant) and “*PDB*” (download PDB coordinates of a mutant). Also, residue-specific aggregation scores are plotted as a function of their sequence position in an interactive graph at the bottom of the “*Automated mutations*” tab. Aggregation propensities of different candidates can be compared by clicking the specific names in the graph’s legend.

**Figure 3.**
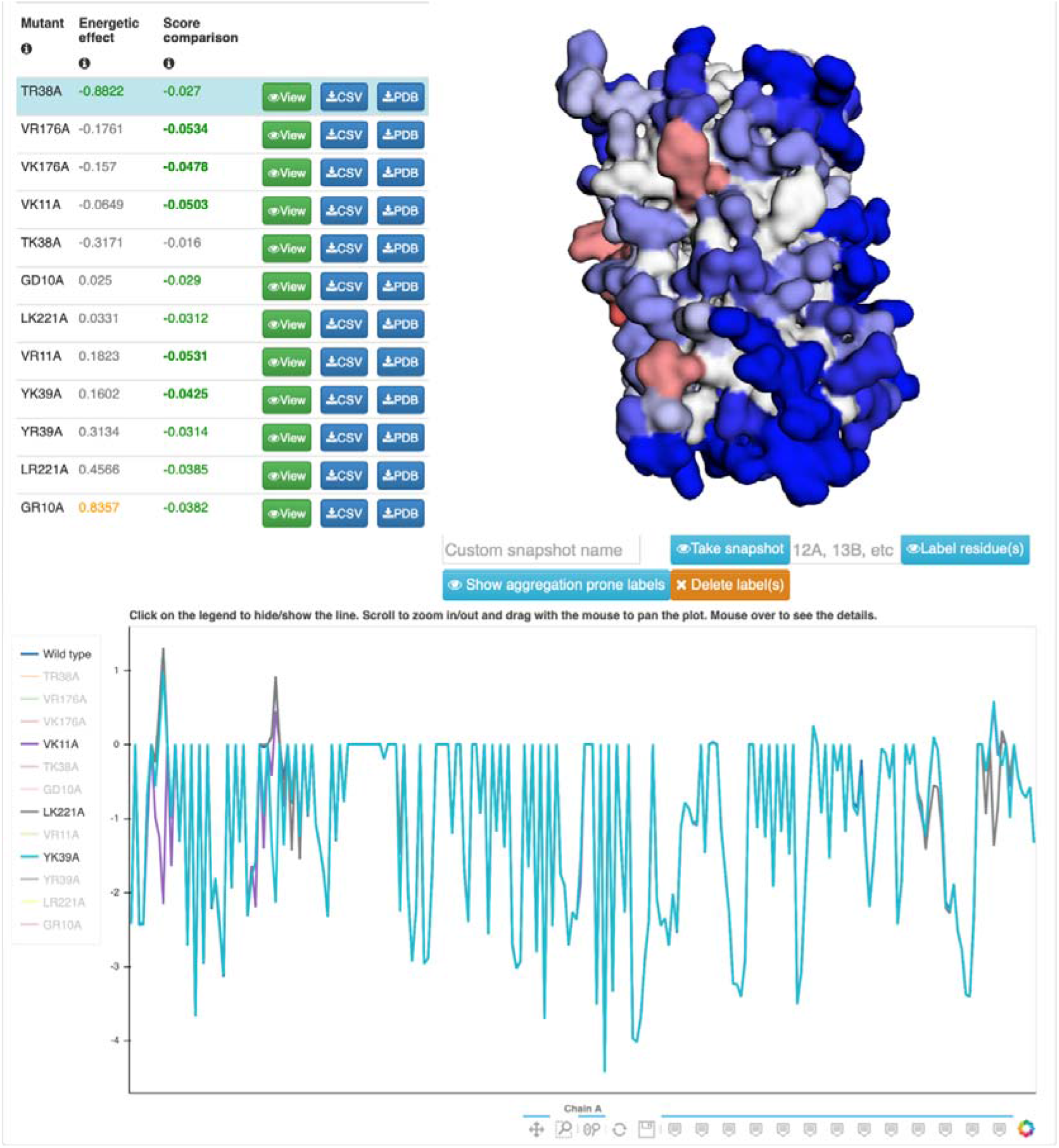
A3D 2.0 automated mutations output for frGFP protein (PDB: 2B3Q). A3D 2.0 suggestions of solubilizing mutant variants are compiled in the table on the left. The table includes information about mutants’ stability and average aggregation scores. The interactive visualization tool (right) allows us to inspect these mutants individually. As well, the specific aggregation propensities of these proposed mutations can be compared in the plot. Here we observe the aggregation profile of the *wild type* frGFP and V11K (purple), Y39K (blue), and L221K (grey) mutants.

In order to facilitate the redesign of protein structures, first, we recommend to perform the exploration of relevant STAPs of the protein under study using static, but also dynamic A3D analysis. This information will be decisive in the selection of mutant candidates and should complement insights on known regions involved in protein function. With this information in hand, we recommend focusing the engineering exercise on potentially harmful regions that are not involved in protein activity. The election should be restricted to those mutants with ΔA3D_score_<0 combined with a neutral or stabilizing effect, always prioritizing ΔΔG<0. Finally, for proteins that exhibit a STAP that can be targeted by different aggregation-prone amino acids, and proteins with two or more APRs, we recommend including a combination of the mutations suggested by the automated mutations tool. An ΔA3D_score_<0 and ΔΔG<0 for those multiple mutants must be verified with the manual mutation tool, before any experimental assay.

### 2.5. Structure visualization and Gallery

A3D 2.0 uses the 3Dmol library for protein structure visualization in an interactive interface. The visualization tool is accessible on the classic *“Structure”* visualization tab and has also been included in *“Dynamic mode details”;* likewise, it is also included in the *“Automated mutations”* algorithm section (Fig 2 and 3). The output structures are colored according to the A3D scale: soluble regions in blue, aggregation-prone regions in red. The color intensity denotes score values: dark-red/ dark-blue indicate the highest/lowest aggregation scores. White and light-red/light-blue depict those amino acids with scores close to 0. In addition to the standard tools to manipulate protein structures, including zooming, structure orientation, downloading of .pdb coordinates, and automatic rotation video, the new version of A3D allows us to manually label specific amino acids or automatically identify those residues with positive aggregation propensity (*see* **Note 11**). Besides, users can always take a snapshot of the visualization, which will be automatically saved on the “*Gallery*” tab, where they will be available as downloadable jpeg image files (*see* **Note 12**).

### 2.6. Example of optimization of protein solubility

Green fluorescence protein (GFP) is a globular protein that emits green fluorescence when exposed to blue light. It was first isolated from the jellyfish *Aqueroa victoria*. It has been extensively employed as a fluorescent reporter to study essential mechanisms of cellular functioning and protein stability. It is one of the most widely used proteins in biological and medical research (the discovery and development of GFP were awarded the 2008 Nobel prize in Chemistry (41)). *Wild type* GFP exhibited poor solubility, which compromised its direct biologic application (42). Since its discovery, multiple studies managed to redesign GFP into modified versions with optimized properties (43). Folding reporter GFP (frGFP) is a variant of the GFP that incorporates the “cycle-3” mutations and the enhanced GFP mutations to the original sequence, which provide increased stability and higher fluorescence brightness (44). A3D 2.0 static runs of the *wild type* GFP (1EMA) and frGFP (2B3Q) report average scores of −0.9342 and −1.0282, respectively. frGFP displays less exposed APRs, which correlates with its observed increased solubility (Fig 4A and 4B). However, a manual inspection of the frGFP’s output reveals the presence of several remaining APRs, clusters of residues with an A3D score >0. “*Aggrescan3D plot*” and “*Aggrescan3D scores*” screens constitute useful interfaces to detect and inspect these residues. Three of these residues – V11, Y39, and V221 – are included in a prominent STAP in frGFP’s surface, yielding A3D values of 1.3585, 0.9007, and 0.3512, respectively. They can be tracked in the “*Structure*” output interface, displayed as red-colored residues (Fig 4B).

**Figure 4.**
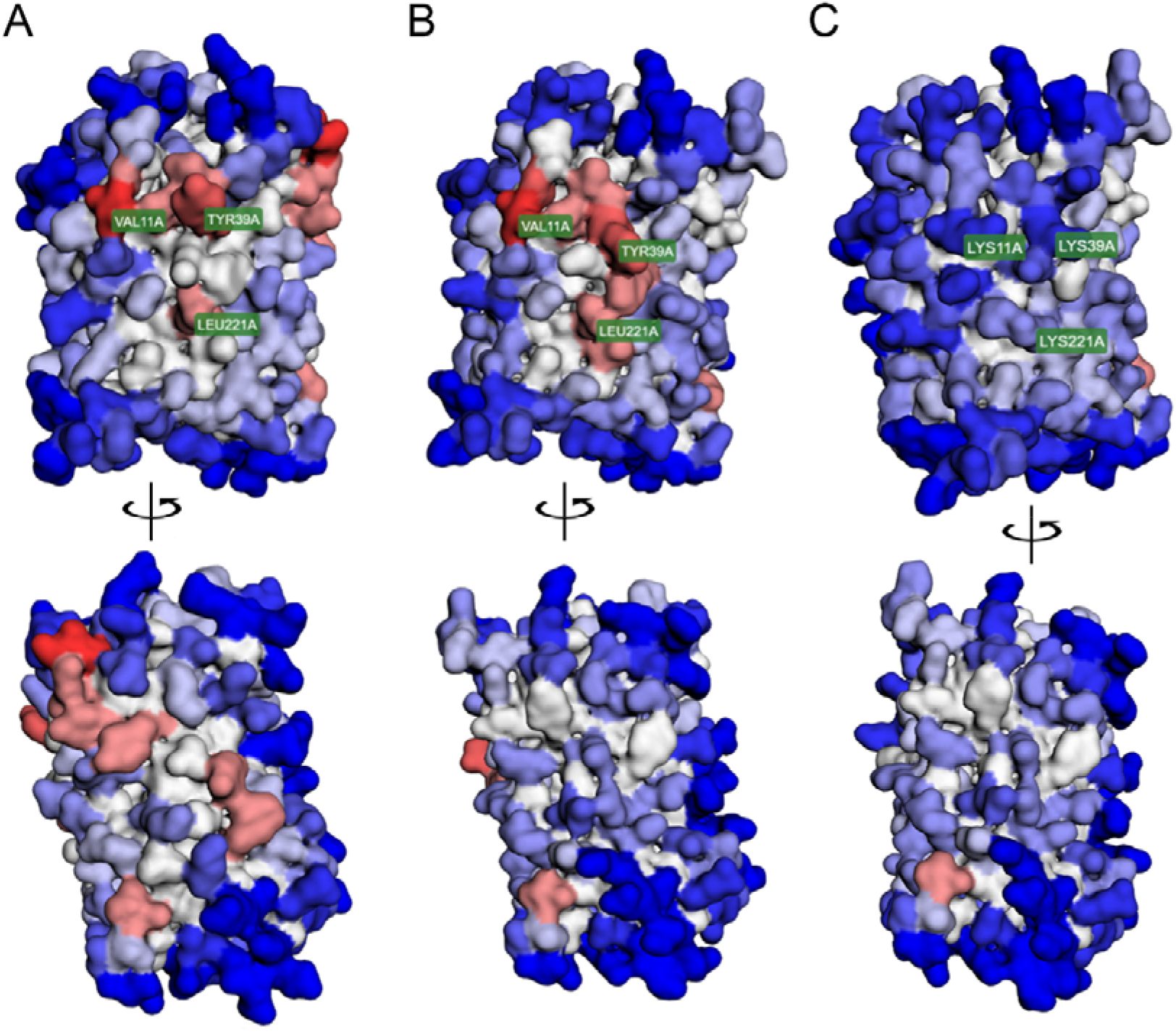
Redesigning an frGFP variant with enhanced solubility. A3D run of the wtGFP (A) and frGFP (B) in static mode; PDBs 1EMA and 2B3Q, respectively. Amino acids are colored according to the aggregation scale of A3D, in which red corresponds to aggregation-prone residues, and blue corresponds to soluble regions. The three residues involved in the frGFP STAP are labeled as V11, Y39, and L221. (C) A3D run of the redesigned frGFP/KKK triple mutant in static mode. Bottom images correspond to a 180° rotation of the structures.

The re-analysis of the frGFP with the enhanced tool renders a list of suggested mutations that might improve the solubility of the protein (Fig 3). The automated mutations tool suggested mutant versions for all the residues involved in the STAP mentioned above: V11, Y39, and V221 (*see* **Note 13**). Specific ΔΔG and ΔA3D_score_ values for these residues are collected in Table 1. According to the automated mutations tool, a mutation to lysine is the best-ranked option for all three residues, although aspartic acid reported an equal or more protective ΔA3D_score_ scores. These data suggest that mutations to positively charged amino acids are better tolerated by the frGFP structure, being thermodynamically more stable. Their impact on frGFP aggregation profiles can be further inspected in the interactive graph of the *automated mutations* output. Mutations to lysine attenuate the aggregation propensity of V11, Y39, and V221, rendering negative A3D scores (Fig 3). V11K, Y39K, and L221K mutations can be combined to abrogate the presence of frGFP’s STAPs completely. The analysis of the frGFP/KKK mutant with the manual mutation tool of A3D 2.0 reports −1.133 average A3D score, an increment of −0.105, and a neutral impact on ΔΔG of 0.017 kcal/mol. In contrast, the frGFP/DDD version reports a similar average A3D score −1.139 but an unfavorable ΔΔG of 3.545 kcal/mol, which would significantly compromise the protein stability. Experimental data support the automated predictions of A3D 2.0 (45). The frGFP/KKK mutant efficiently folds into the native conformation and possess stability similar to that of the *wild type* frGFP. However, it is significantly more resistant to aggregation.

**Table 1.**
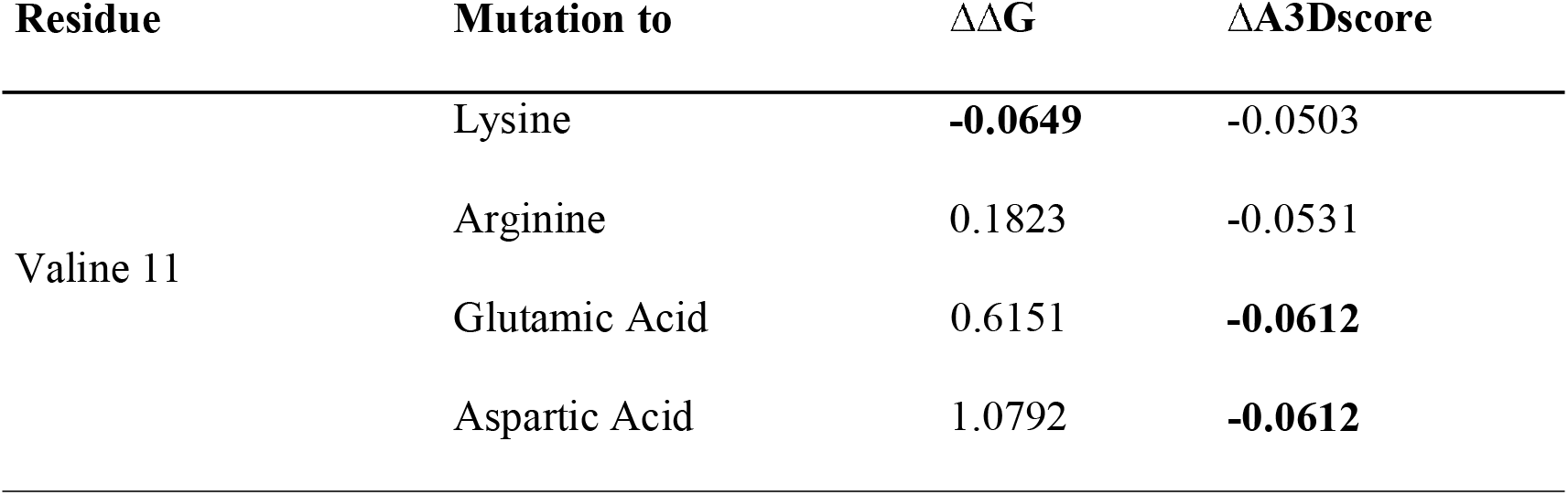
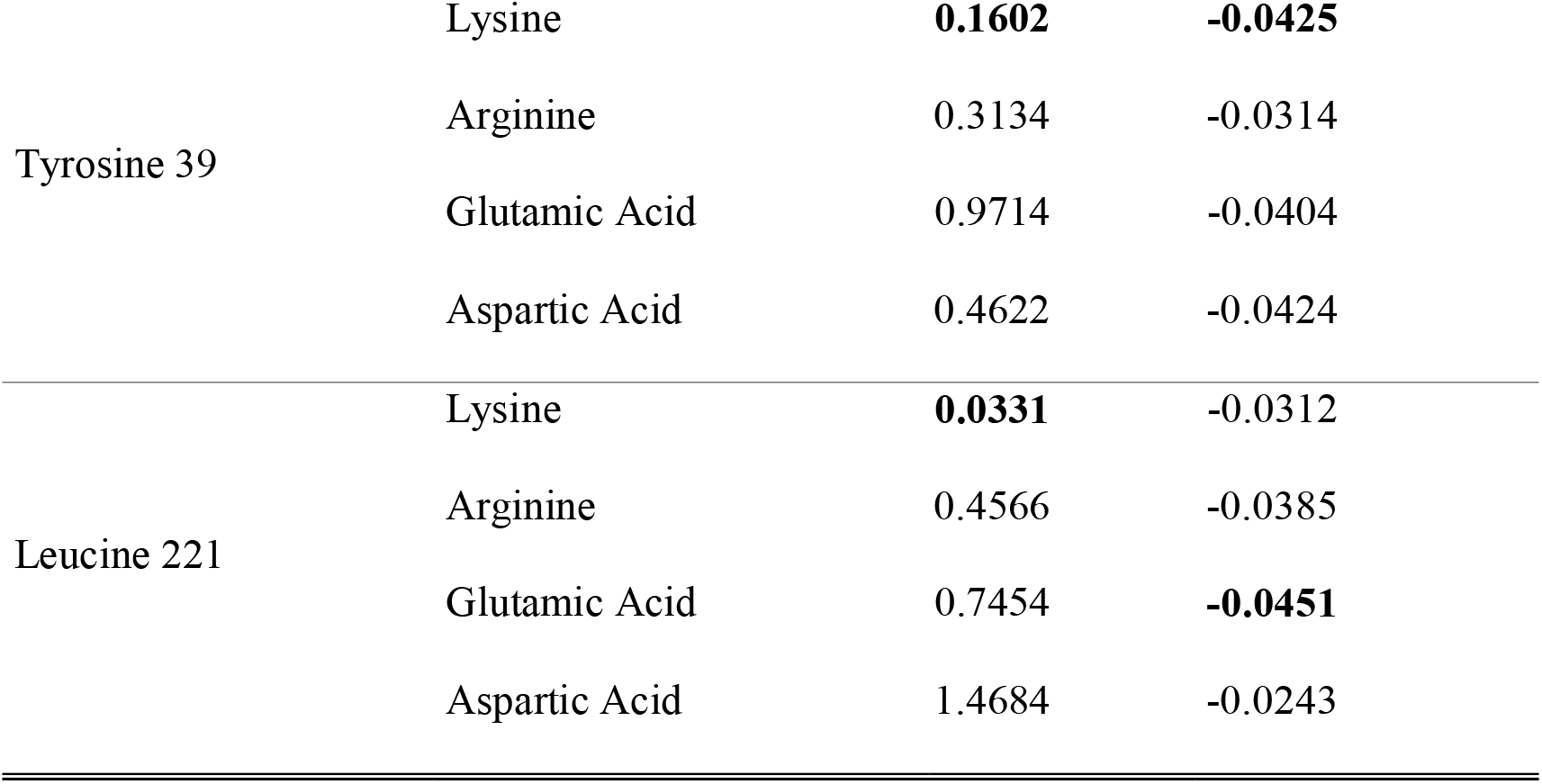
Ranking of the solubilizing mutations suggested by the enhanced mutations tool of A3D 2.0 for the residues 11, 39 and 221 of frGFP. The most solubilizing and less destabilizing change for each position is indicated in bold. A successful redesign should take into account both variables.

| Residue | Mutation to | $\Delta\Delta G$ | $\Delta A3D_{\text{score}}$ |
| --- | --- | --- | --- |
| Valine 11 | Lysine | **-0.0649** | -0.0503 |
|  | Arginine | 0.1823 | -0.0531 |
|  | Glutamic Acid | 0.6151 | **-0.0612** |
|  | Aspartic Acid | 1.0792 | **-0.0612** |
| Tyrosine 39 | Lysine | **0.1602** | **-0.0425** |
|  | Arginine | 0.3134 | -0.0314 |
|  | Glutamic Acid | 0.9714 | -0.0404 |
|  | Aspartic Acid | 0.4622 | -0.0424 |
| Leucine 221 | Lysine | **0.0331** | -0.0312 |
|  | Arginine | 0.4566 | -0.0385 |
|  | Glutamic Acid | 0.7454 | **-0.0451** |
|  | Aspartic Acid | 1.4684 | -0.0243 |

### 2.7. A3D 2.0 RESTful Service

A3D 2.0 also implements a RESTful application program interface (API) that allows full server access through the command-line. In this section, we briefly review A3D 2.0 RESTful service to provide a comprehensible guide to easily automatize A3D 2.0 usage without needing much experience in programming. The RESTful module is more suitable for the analysis of extensive collections of proteins or mutants, skipping the analysis of case-by-case inputs in the webpage and tedious data collection. Additionally, the A3D 2.0 RESTful service allows the integration of the A3D 2.0 functionality into alternative bioinformatics pipelines.

To submit a job for a selected protein, (in this example PDB 1PHT) with default parameters, run the following Python script (*see* **Note 14**):

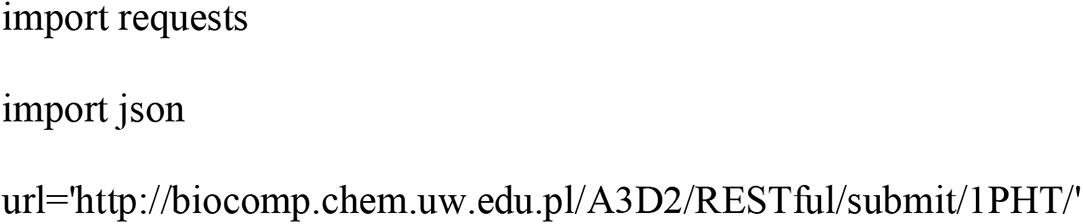

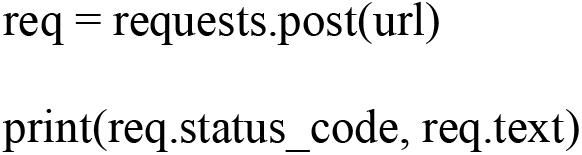

A3D RESTful mode allows access to all modes of A3D by passing a json object with optional information: establish a project name, notify users at a given email, select the distance used for calculation, mutate residues or perform automatic mutations described previously in the front page section, and an option to disable Foldx energy minimization before running the prediction (*see* **Note 15**. The following script is an example, in which we manually mutate two glycines – in positions 1 and 2 – for alanines and run A3D in dynamic mode.

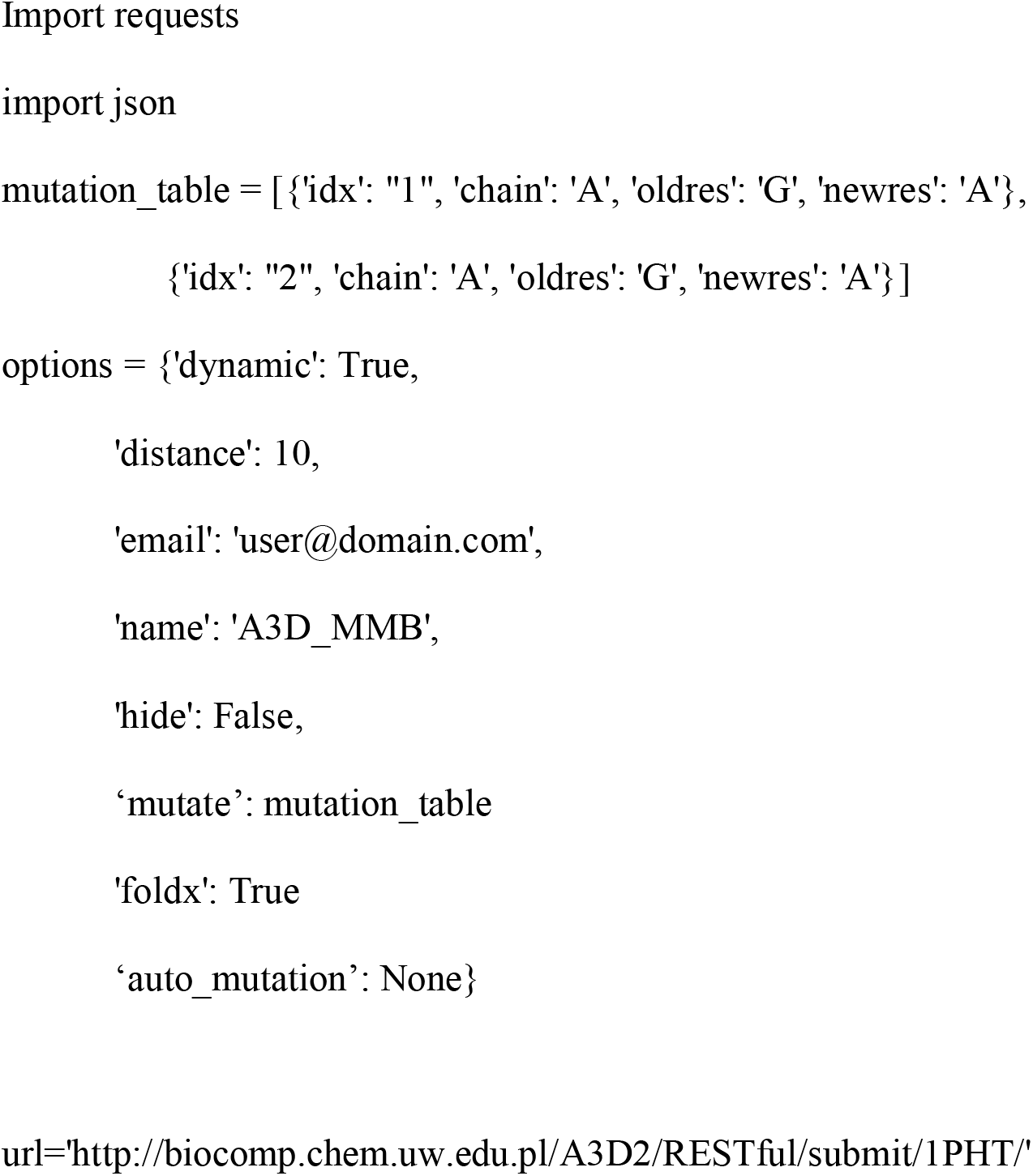

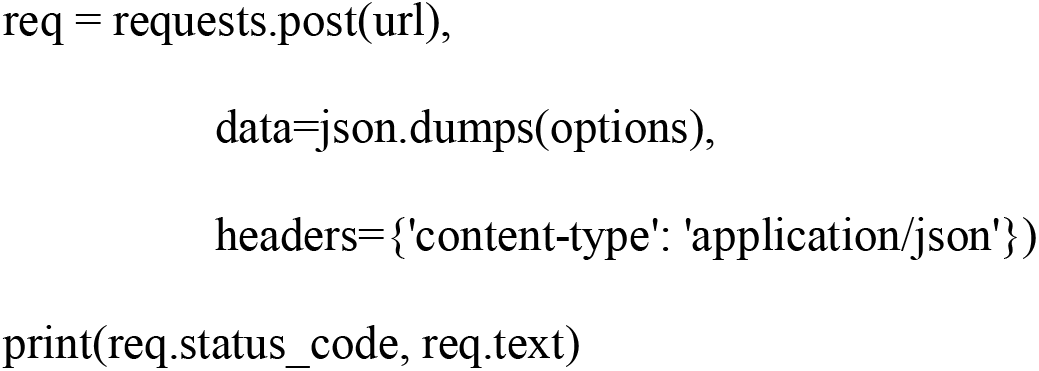

The execution of these scripts prints a status code and the given job identifier as “jobid” (*see* **Note 16**). Searching this unique job identifier on the A3D web server retrieves the standard online output. The analysis can also be acquired as a JSON file containing numerical results. We can systematically obtain the A3D average score for hypothetical jobid ‘jobid1’ with the following script (*see* **Note 17**):

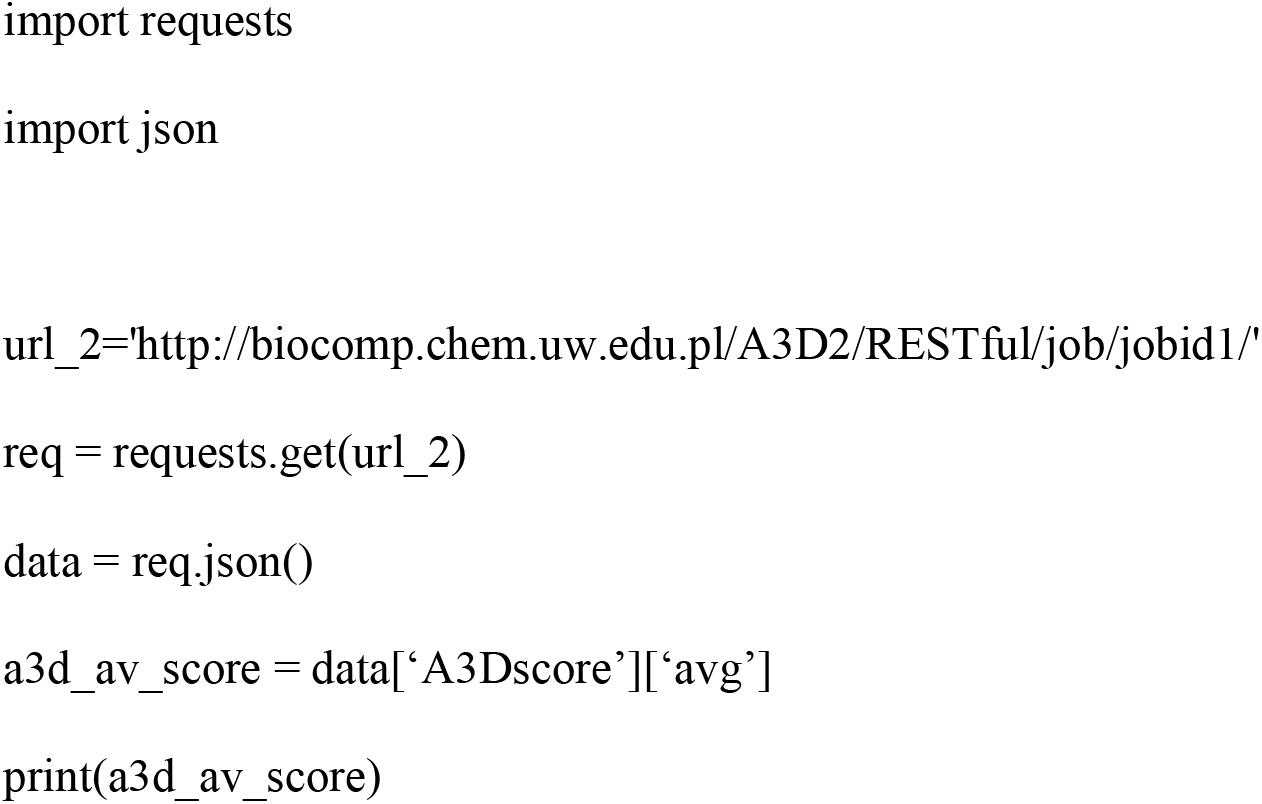

With this tool, users can automatize predictions for a list of structures. In the following example, we show how to use a file with a list of PDB structure identifiers named list.txt and submit them to A3D via the RESTful mode (*see* **Note 18**). Users should copy-paste this code into a text file and save it as submit_pdbsMMB.py. Then, open the terminal in macOS or Linux or command line in Windows and change directory by typing ‘cd’ and the specific path to where the file is located.

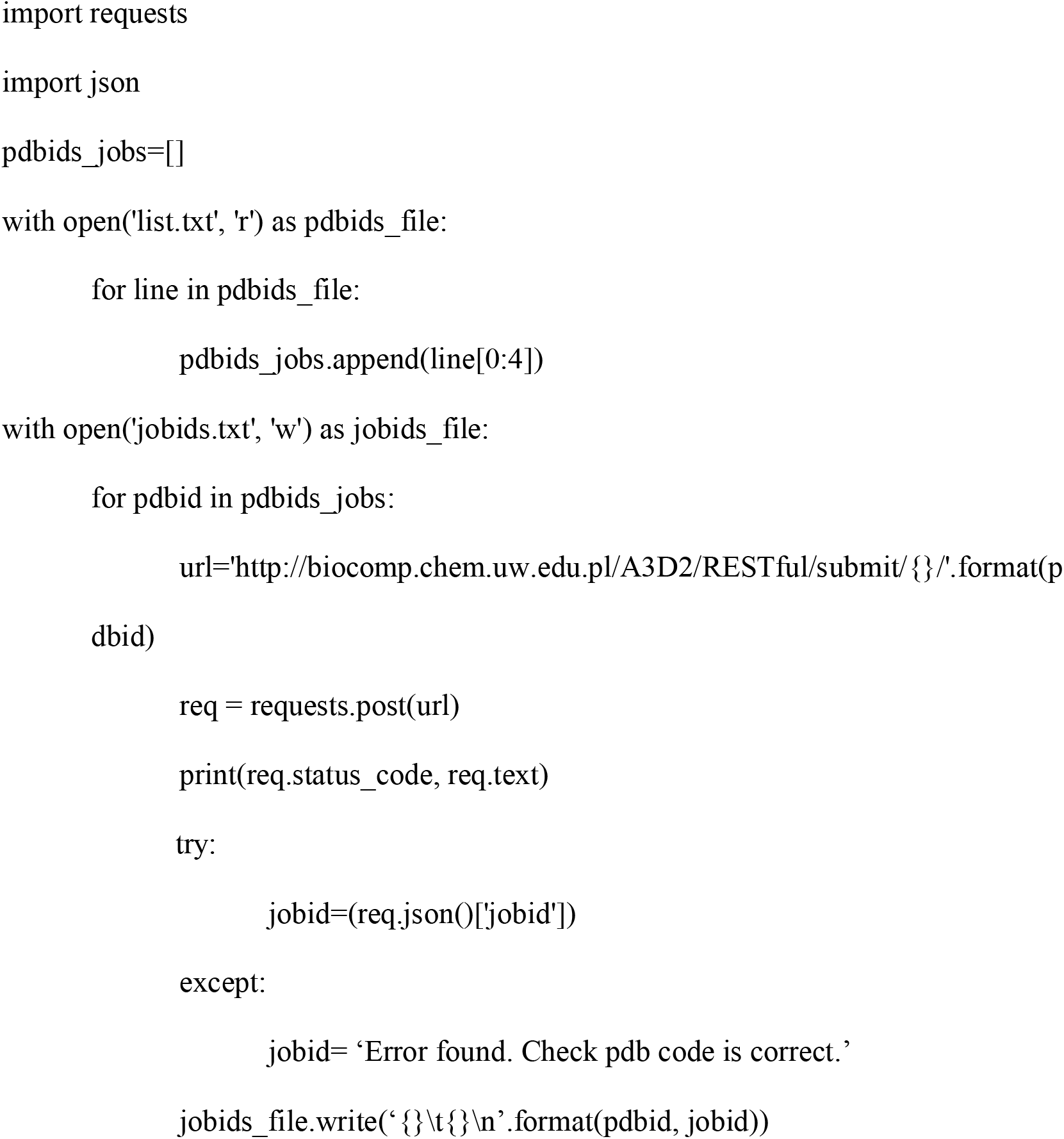

The previous code will retrieve a new file called ‘jobids.txt’ that include a list of pdbs and their job identifiers. Once A3D 2.0 has processed the jobs, their average scores can be compared by executing the following script (*see* **Note 19**):

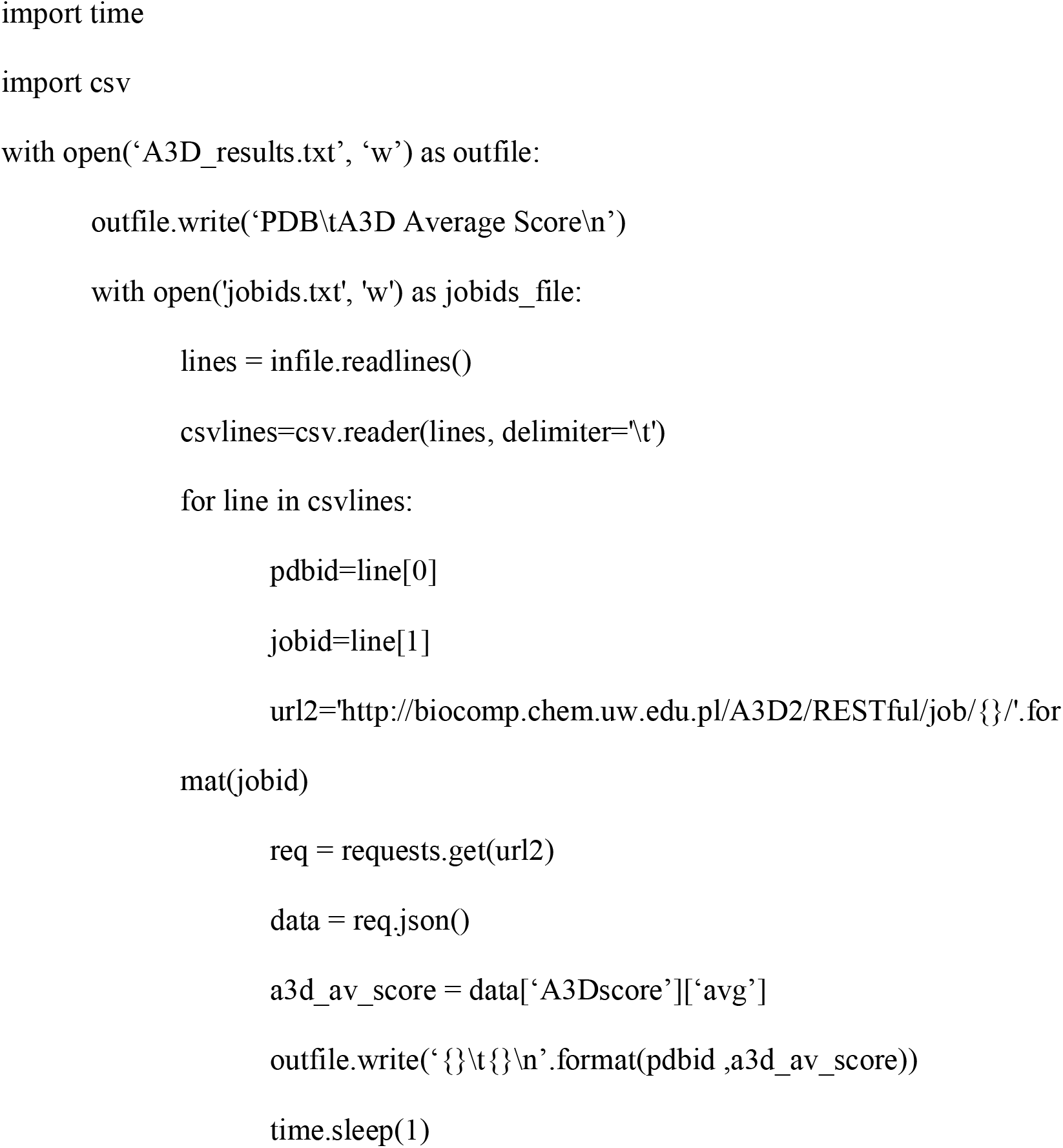

This will generate a new file, A3D_results.txt, with the PDB identifier and A3D average score in adjacent columns. Users can find different examples of how to interact with A3D RESTful mode in https://github.com/PPMC-lab/A3Dscripts.

## 3. Concluding remark

The interplay between protein function and aggregation entails the unavoidable presence of APRs in protein sequences and structures. This endorses a vast majority of proteins with an inherent propensity to precipitate, especially when out of their biological context. Since the outbreak of the first generation of linear algorithms (using sequence information only) 15 years ago, new computational tools for the prediction of protein aggregation have evolved hand-in-hand with the advances in the understanding of the aggregation process. These versatile computational strategies have demonstrated their potency to spot these dangerous regions in protein structures and proteomes, although their implementation in protein engineering routines is still scarce.

The A3D 2.0 tool introduces considerable improvements that try to address current needs by extending the applications of dynamic mode predictions, including the contribution of structure stability and incorporating an automatic tool to enhance protein solubility. This set of upgraded features harmonizes with the server’s predictions and visualization. Overall, A3D 2.0 constitutes a powerful routine to model protein aggregation and solubility in a fast and easy-to-use interface adapted for non-expert users.

## 4. Notes

1. Stability calculations always consider the input structure as a reference to determine free energy increments. It is crucial to take this into account if we perform a series of mutations and their joint analysis or model a protein variant back to its wild type form.
2. The accuracy of FoldX predictions depends on the quality of the protein crystal and the completeness of the PDB structure (28).
3. Input PDB structures for a dynamic mode analysis require the complete set of backbone atoms. Partially solved structures are not suitable.
4. In order to assist in data analysis, we recommend adjusting the X-axis to protein’s length and zoom out the Y-axis for fast analysis of the aggregation profile (place the cursor on the specific axis and scroll up/down). This configuration facilitates the identification of regions or specific amino acids with differential aggregation propensities. Once they are identified, zoom in or use the crop tool to scrutinize that region further and determine the specific amino acids involved in the STAP. If needed, graphs can be captured at any time and downloaded as snapshots by clicking the floppy disk icon.
5. RMSF (Root Mean Square Fluctuation) here is a measure of the deviation of the position of a residue *i* (C-alpha atom) with respect to its reference position over CABS-flex simulation time, expressed in Ångstroms. Therefore, RMSFs plots as a function of residue number illustrate chain flexibility.
6. CABS-flex models are ranked according to their average A3D score. Accordingly, models containing informative STAPs might be considered low-ranking because of a higher global solubility, and vice versa, top-ranking models might not possess any prominent STAP but a significant average aggregation propensity. We firmly recommend exploring the interactive tools provided in the “dynamics” output and manually curate all CABS-flex simulations.
7. Mutations tool and enhance solubility (automated mutations) tool are incompatible. If they are both selected, A3D 2.0 will only execute the mutations tool. We recommend starting with an initial automated mutation analysis to obtain solubilizing candidates and proceed to the manual inspection and optimized modifications with the mutations tool.
8. For some proteins, relevant aggregation-prone regions can only be detected in dynamic runs of A3D. In these cases, users should download the PDB structure of the specific CABS-flex model that contains the aggregation-prone region and use it as the initial structure of the enhanced solubility (automated mutations) tool.
9. Charged residues usually flank stretches of apolar residues in proteins sequences and structures and attenuate local hydrophobicity, functioning as aggregation gatekeepers (46).
10. Some particular proteins require the presence of exposed hydrophobic residues to perform their biologic function. This is the case of the complementarity-determining regions (CDR) of antibodies, which usually encompass certain hydrophobicity to recognize and interact with their targets. Despite being beneficial for protein solubility, modifying CDR residues might compromise antigen recognition seriously.
11. To label a residue manually, enter the residue number followed by the chain letter code into the “label residue” box (for example, 24A). Combining the manual label tool with both the information obtained in the “Aggrescan3D plot” and “Aggrescan3D score” will help to place STAPs in the protein structure.
12. We encourage capturing snapshots while analyzing protein structures. Remember to name the snapshots in the “take a snapshot” box before capturing the image.
13. Additional top-scoring mutations were reported for the threonine 38 and valine 176. Inspection of these amino acids allowed to conclude they do not suit the redesign of frGFP. First, T38R top-ranking mutation does not come from a protective ΔA3Dscore, but an extreme stabilization of the protein structure. Second, V176R and V176K protective ΔA3Dscore do not stem from the attenuation of a potentially dangerous APR, but from a moderate solubilizing effect on a more extended neutral region.
14. To run another PDB, change the 1PHT in the URL for the desired PDB code. Further options, including single chain or locally stored PDB, can be found in the server tutorial.
15. The output screen can provide different codes: 200 means the job has been posted correctly, while 400 or 404 are different errors.
16. For any of the fields, A3D 2.0 will apply default parameters unless otherwise specified.
17. Users should introduce the jobID obtained in the previous step in the url_2, instead of the jobid1 code used in the example.
18. The list.txt file must have one PDB identifier per line without a header and be saved in the same folder as the script.
19. Both the script and jobids.txt files should be saved in the same folder.

## Acknowledgements

This work was funded by the Spanish Ministry of Economy and Competitiveness BIO2016-78310-R to S.V and by ICREA, ICREA-Academia 2015 to S.V. J. S. and J.P. were supported by the Spanish Ministry of Science and Innovation via a doctoral grant (FPU17/01157 and FPU14/07161) . Conflict of Interest: none declared.

